# Mapping human social brain specialisation beyond the neuron using multimodal imaging in human infants

**DOI:** 10.1101/2022.11.08.514512

**Authors:** Maheen Siddiqui, Paola Pinti, Sabrina Brigadoi, Sarah Lloyd-Fox, Clare E. Elwell, Mark H. Johnson, Ilias Tachtsidis, Emily J.H. Jones

## Abstract

The specialised regional functionality of the mature human cortex partly emerges through experience-dependent specialisation during early development. Our existing understanding of this process is based on evidence from unitary imaging modalities and has thus focused on isolated changes in spatial or temporal precision of neural or haemodynamic activation alone, giving an incomplete picture of the process. We speculate that neural specialisation of function will be underpinned by better coordinated haemodynamic and metabolic changes in a broader orchestrated physiological response. Thus, we present a harmonised framework in which specialisation is indexed by the emergence of coupling between neuronal activity and vascular supply of oxygen and energy. Here, we combine simultaneous measures of coordinated neural activity (EEG), metabolic rate and oxygenated blood supply (broadband near-infrared spectroscopy) to measure emerging specialisation in the infant brain. In 4-to-7-month-old infants, we show that social processing is accompanied by spatially and temporally specific increases in coupled activation in the temporal-parietal junction, a core hub region of the adult social brain. During non-social processing coupled activation decreased in the same region, indicating specificity to social processing. Coupling was strongest with high frequency brain activity (beta and gamma), consistent with the greater energetic requirements and more localised action of high frequency brain activity. We conclude that functional specialisation of the brain is a coordinated activity across neural, haemodynamic, and metabolic changes, and our ability to measure these simultaneously opens new vistas in understanding how the brain is shaped by its environment.

## Introduction

The adult brain is highly specialised, with core networks coordinating to subserve complex behaviours. This specialised functioning emerges across development through a combination of genetically influenced brain architecture and experience-dependent and experience-expectant learning processes (*1*). This interaction between predisposition and change with experience has been closely studied in the domain of social interaction, where neonates attended preferentially to faces (*2*) but expertise in recognition, communication, and initiation emerge gradually over time (*1, 3*). Social communication is core to human interaction, and our ability to live in extended-family groups has been linked to the evolution of advanced cognitive abilities (*4*). Thus, understanding the processes that shape social brain development is critical to understanding the ontogeny and phylogeny of our species.

In adulthood, social interaction is partially subserved by a network of specialised regions that include the amygdala, fusiform gyrus, superior temporal sulcus, and medial prefrontal cortex (*5*). However, the mechanisms through which this network becomes specialised for social processing remains unclear, in part because studies have typically used single modalities sensitive to distinct aspects of brain function. For example, the N170 event-related electroencephalographic brain response indexes expertise with faces and can be sourced to the fusiform gyrus (*6*). This response can be detected by 4 months (*7*), but its sensitivity to configural processing develops over the first year of life (*8*). Functional magnetic resonance imaging (fMRI) indicates that core regions of the social brain (particular the fusiform face area) show increases in oxygenated haemoglobin delivery in response to faces by 4-9 months (*9*). Functional near-infrared spectroscopy (fNIRS) studies show that oxygenated haemoglobin delivery in response to naturalistic social videos in a broad region of temporal cortex emerges over the first hours of life (*10*). Thus, work with single modalities indicates experience-dependent changes in specialised brain activity across the first year of life but does not yield insights into the underpinning mechanisms.

Interactive specialisation is a theory of brain development that posits that competition between brain regions for acquiring function drives specialisation (*3*). This can be indexed through a reduction in the spatial extent of neural (and vascular) responses to a particular stimulus category and a concomitant increase in selectivity in responsive regions (*11*). One mechanism that could underpin this competition is the limited energetic resources available to the infant brain. The brain is an energetically costly organ, consuming 20-25% of the body’s energy in adulthood while representing only 2% of the body’s mass (*12, 13*). There are also substantial developmental changes in the brain’s energy consumption; in the first year of life, up to 60% of available energy is used by the brain (*14*). When brain regions become functionally active (for example during stimulus processing) neurons fire more rapidly, requiring greater supplies of adenosine triphosphate or ATP (energy stores). Producing ATP requires oxygen, and this is supplied through a localised increase in oxygenated haemoglobin in the blood. Increases in oxygenated haemoglobin do not happen concurrently in all brain areas, and there are spatial dependencies between activated and deactivated regions in the adult brain (*15*). Energy supplies are important to synaptic plasticity, memory and learning (*16*), and the mechanism through which energy supplies are coupled to activation (neurovascular coupling) also develops through experience-dependent specialisation in the infant brain (*17*). Thus, we propose that examining the coupling between neuronal activity and energy supply will provide the most sensitive measure of the emergence of specialised brain function in the infant brain.

Broadband near-infrared spectroscopy (or bNIRS) is a new technique that can be used to quantify the relationship between the neuronal, hemodynamic, and metabolic activity in the infants’ brain as it allows the simultaneous and non-invasive acquisition of haemodynamic and metabolic activity concurrently with EEG during functional activation. This technology uses a broad range of optical wavelengths which allows the measurement of the oxidation state of mitochondrial respiratory chain enzyme cytochrome-c-oxidase (CCO), thereby providing a direct measure of cellular energy metabolism (*18*). CCO is located in the inner mitochondrial membrane and serves as the terminal electron acceptor in the electron transport chain (ETC). It therefore accounts for 95% of cellular oxygen metabolism. In this way, bNIRS allows non-invasive measurement of cellular energy metabolism alongside haemodynamics/oxygenation in awake infants. We recently showed the feasibility of using bNIRS in 4-to-7-month-old typically developing infants (*19*) and demonstrated the presence of unique task-relevant, regionally specific functional networks where high levels of haemodynamic and metabolic coupling were observed. Here, we integrate this methodology with EEG to identify markers of early brain specialisation with coordinated energetic coupling and neural activity. We develop a novel analysis pipeline to identify localised coupling responses that are modulated by naturalistic social content. We predicted that coupling would be most pronounced in the high-frequency beta and gamma band (*20*–*25*) (*26*), and we hypothesised that we would identify core localised social brain regions with coordinated increases in coupled neural activity, metabolic changes and neurovascular response in the infant brain.

## Results

### Naturalistic social stimuli elicit expected increases in broadband EEG activity

5-month-old infants n=42) viewed naturalistic social and non-social stimuli (Fig 1a) while we concurrently measured EEG and broadband NIRS. Fourier-transform of continuously recorded EEG data from 32 channels (n=35) in one-second segments across the time course of stimulus presentation confirmed robust broadband increases in neural activity in response to social versus non-social stimuli (Fig 1b, replicating (*11*)).

**Figure 1:**
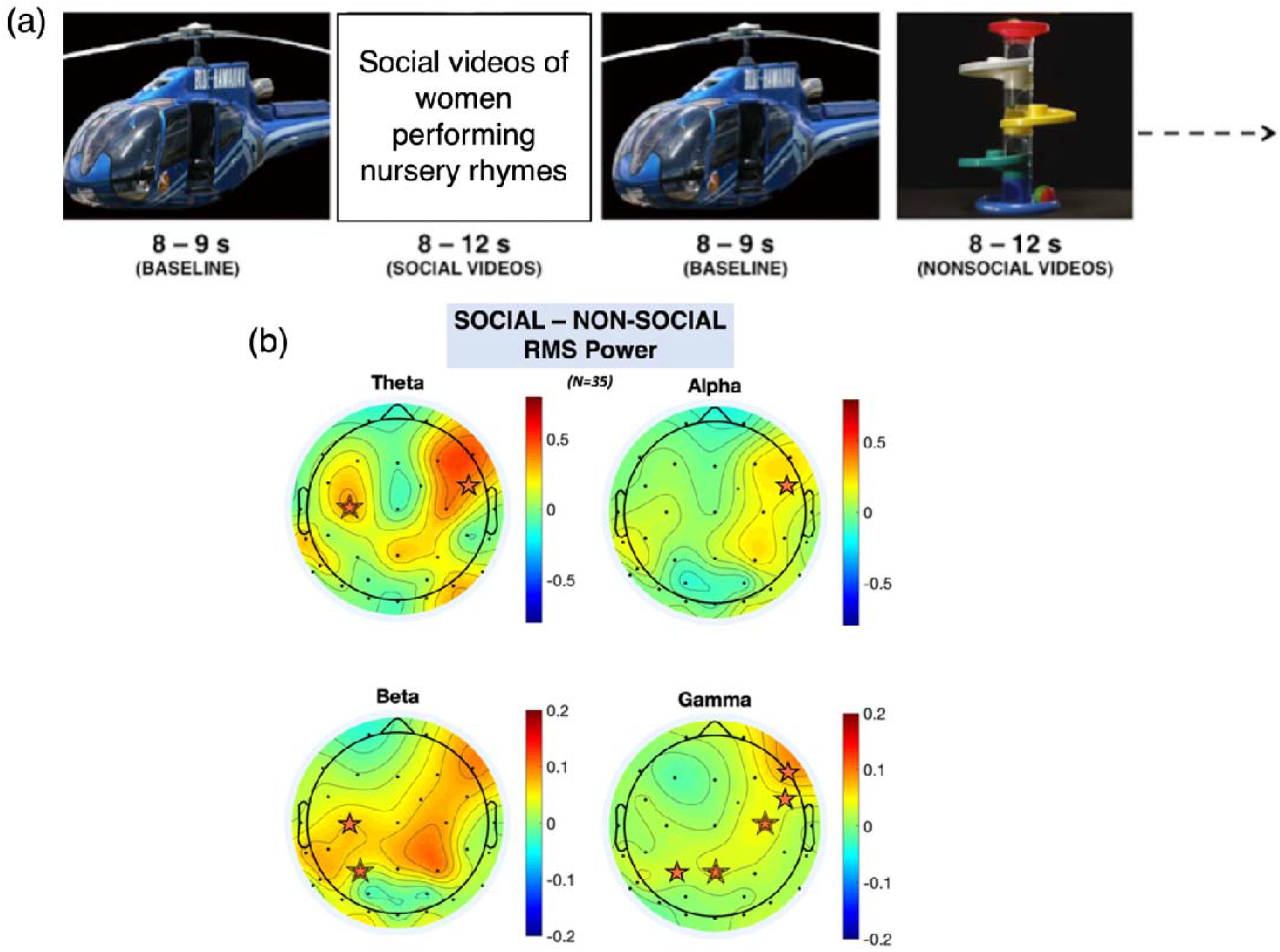
a) Illustration of the paradigm; b) Scalp topographies of the grand average RMS power for theta, alpha, beta, and gamma frequency bands (averaged across participants, averaged across the stimulus period) for the social minus non-social condition. The orange stars indicate statistically significant EEG electrodes where an increase in activity was observed (e.g., increase in response to the social condition compared to the non-social condition) while the grey stars indicate statistically significant EEG electrodes where a decrease in activity was observed; a double line indicates significance after FDR correction.

### Haemodynamic and metabolic coupling and oscillatory activity spatially overlap

A validated method Fig 2f (*27*) applied to the bNIRS data (n=25) identified regions with coupled increases in metabolic function and oxygenated blood flow (*19*). This revealed distinct locations sensitive to social (Fig 2b) and non-social (Fig 2d) processing; the topography of these locations is strikingly similar to the topography of differentiated broadband EEG activity (Fig 2a, c, e).

**Figure 2:**
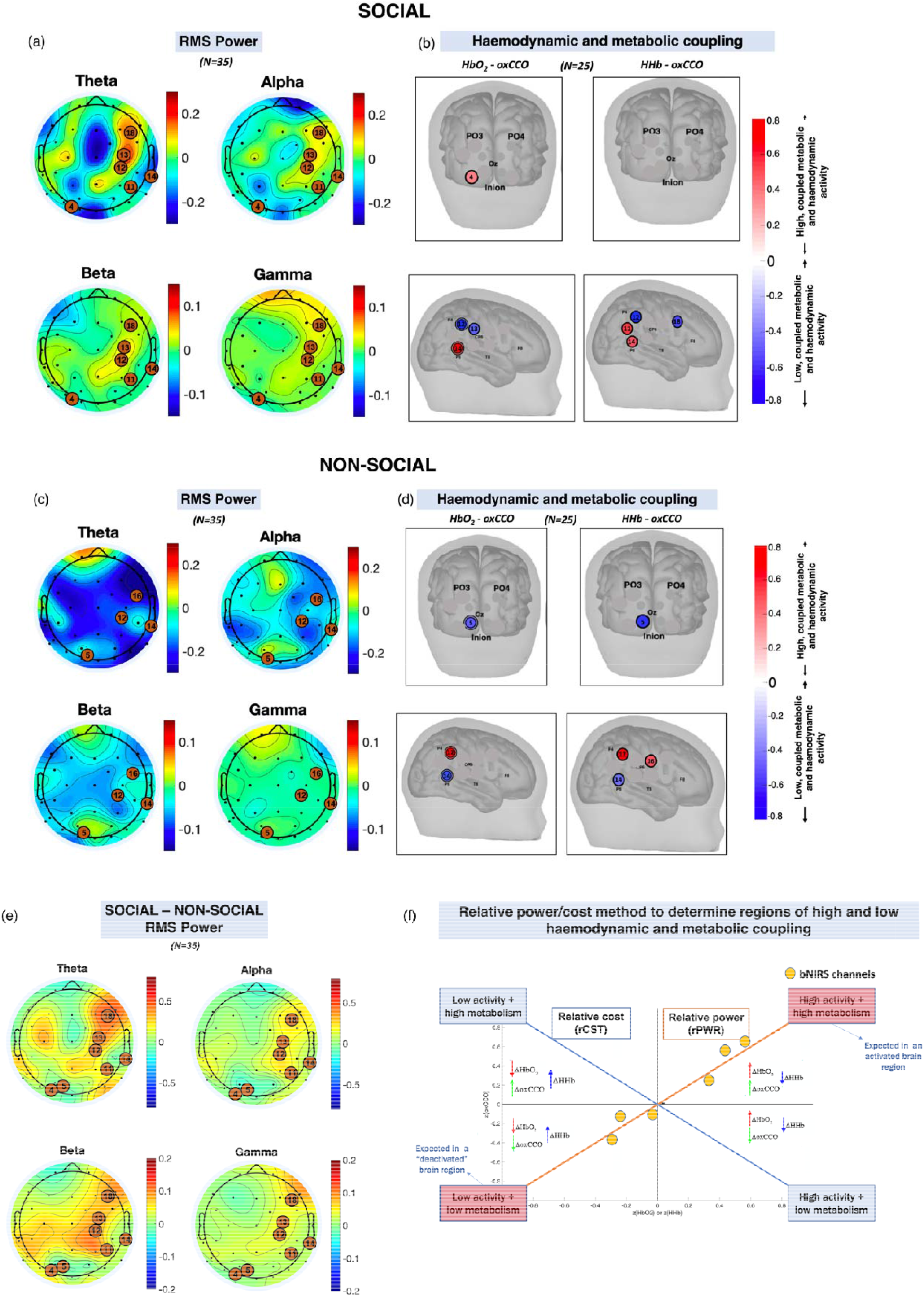
Scalp topographies of the grand average RMS power for theta, alpha, beta, and gamma frequency bands (averaged across participants, averaged across the stimulus period for (a) social and (c) non-social conditions. Locations of high haemodynamic and metabolic coupling for (b) social and (d) non-social condition obtained using (f) the relative power and cost method described in(27, 28).

### Coupled signals highlight specialised activation in the temporal parietal junction

We then convolved the time-course of the within-hemisphere EEG responses with an infant-specific haemodynamic response function (n=17; Fig 3a). A general linear model (GLM) approach was then used to identify FDR-corrected associations between EEG channels and bNIRS channels that showed significant coupling between metabolic response and oxygenated haemoglobin delivery (Fig 2 b, d). We were looking for bNIRS channels showing the expected patterns of positive associations between EEG and oxCCO and HbO_2_ and negative associations with HHb. Figure 3 shows that these associations were primarily concentrated in the beta and gamma bands as predicted (Fig 2 in the supplementary material shows the associations for the theta and alpha bands). Coupled activity was localised to a bNIRS channel (channel 14) positioned over the superior temporal sulcus - temporo-parietal junction region. At this channel, a coupled increase for the social condition and a coupled decrease for the non-social condition was observed (Fig 3 b, c).

**Figure 3:**
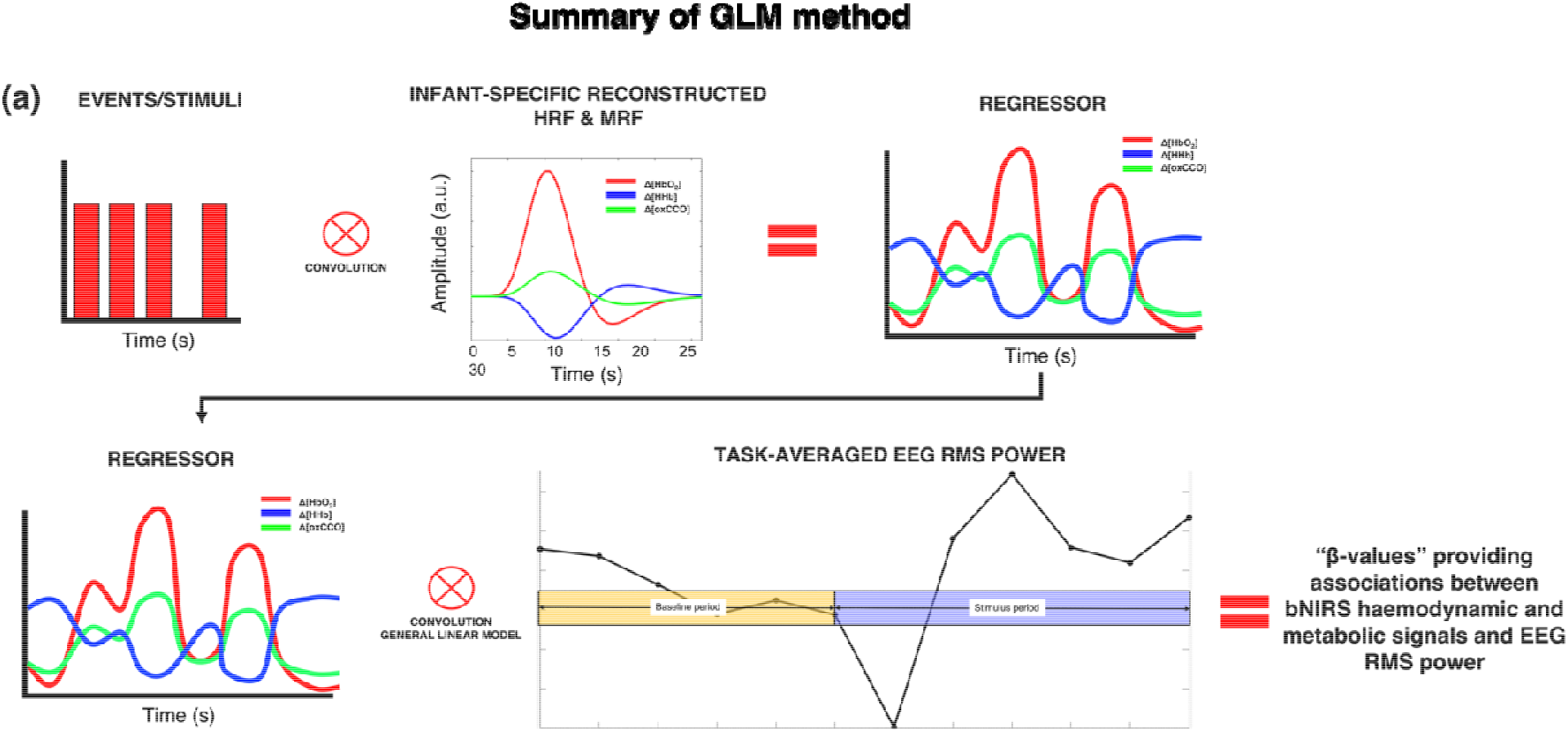
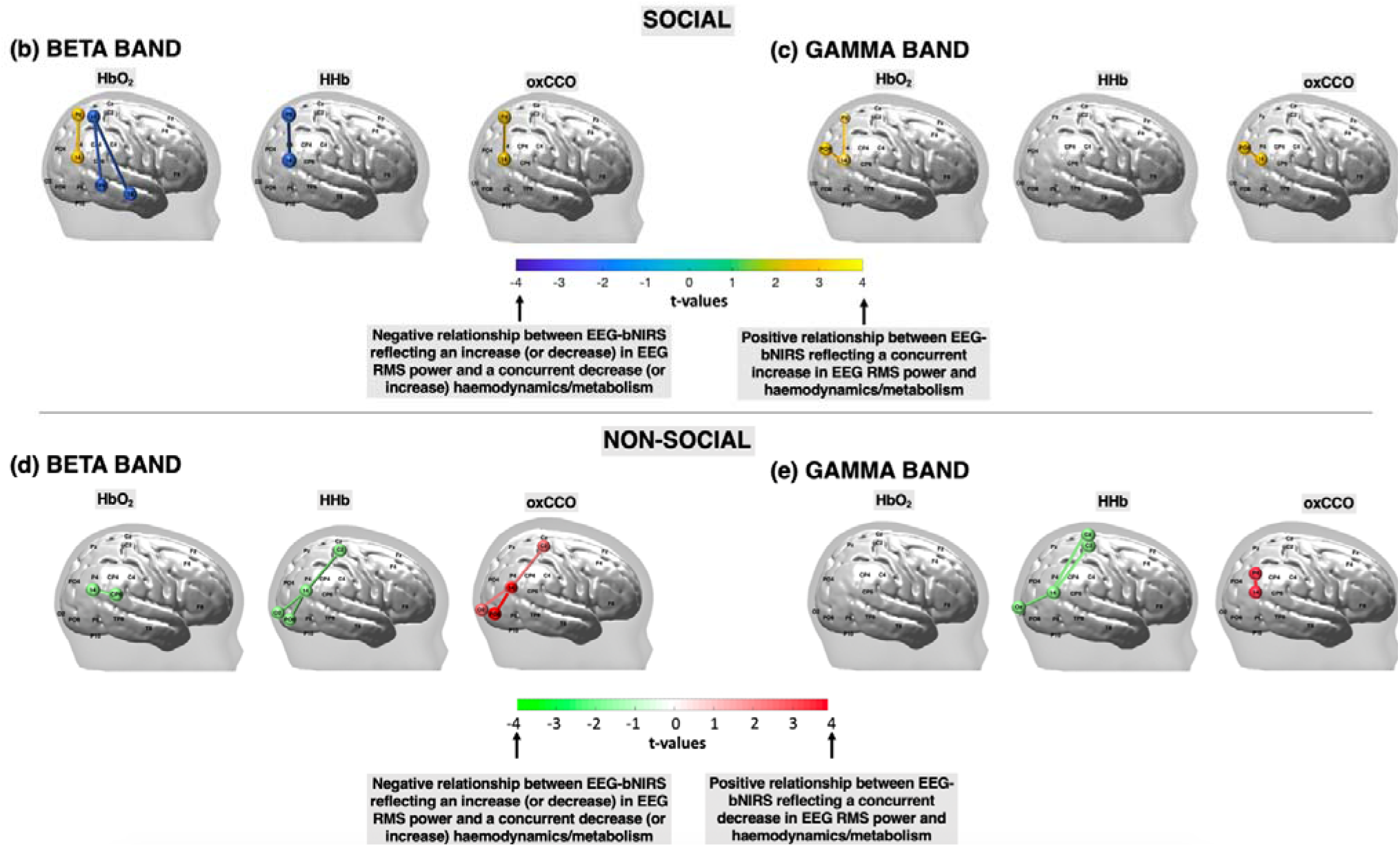
(a) Summary of the procedure for obtaining the associations between bNIRS signals and EEG RMS power at each bNIRS channel combination, for each frequency band. FDR-corrected significant connections between bNIRS channels and EEG electrodes for the beta and gamma bands for the social condition (b-c) and the non-social condition (d-e) for HbO_2_, HHb, and oxCCO. The colour bar represents the t-values from the GLM analysis with a positive t-value representing a significant, positive connection between the bNIRS channel and EEG electrode while a negative t-value represents a negative connection.

Using image reconstruction on the bNIRS data, the spatial sensitivity of the bNIRS location of interest (channel 14) is shown in Figure 4. The method for image reconstruction has been described in detail in the methods section. This indicates that coupled activity was most consistent with the spatial extent of changes in metabolic activity (CCO) and was differentially modulated in the social and non-social conditions.

**Figure 4:**
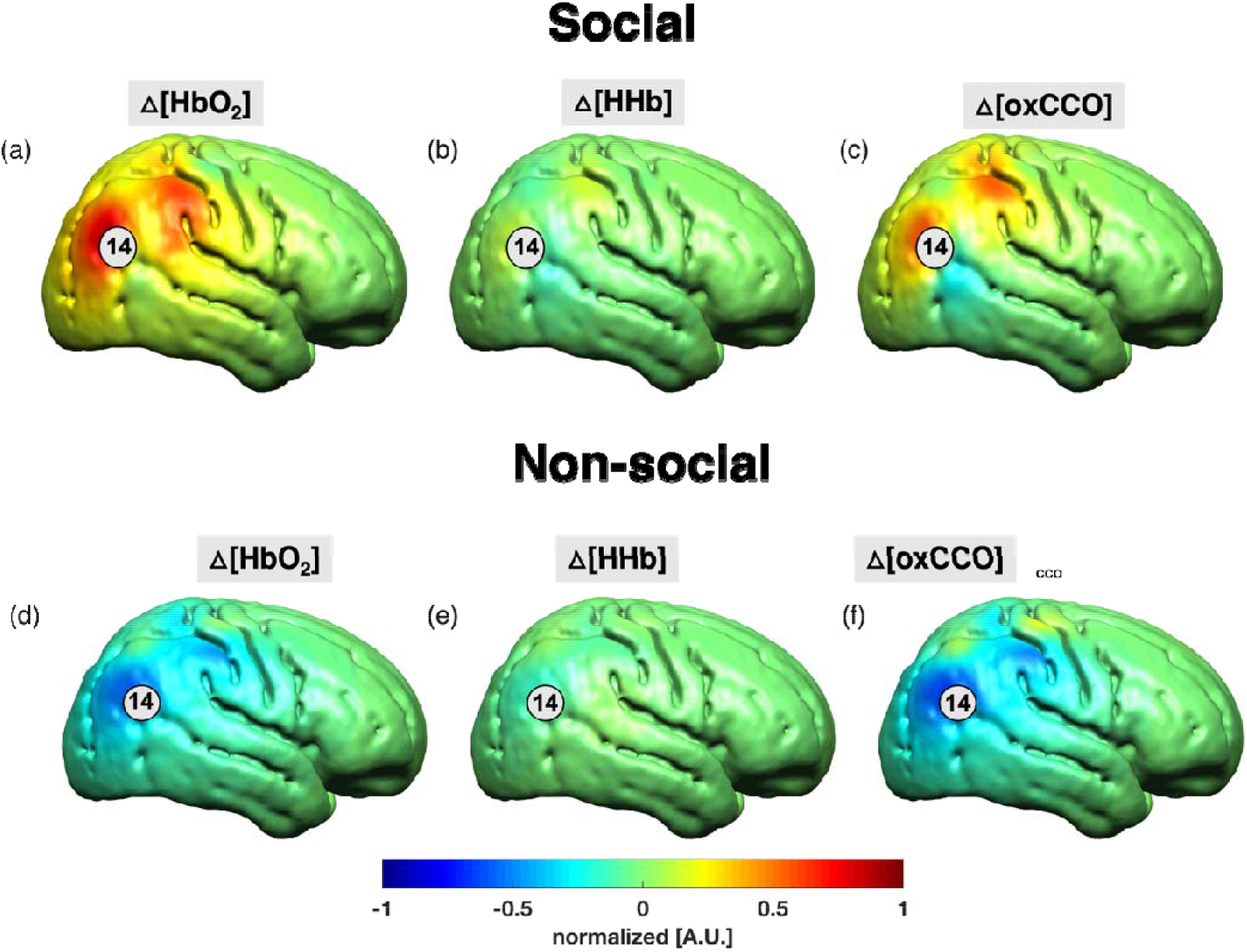
Grand-average image reconstruction at 18 s post-stimulus onset for the social condition (a – c) and the non-social condition (d – f) at a single time point of 18 s post-stimulus onset. The concentration changes for HbO_2_ and HHb were normalised to the maximum concentration change of HbO_2_ while ΔoxCCO was normalised to its own maximum change in concentration. Channel 14 has been indicated.

## Discussion

We conducted a multimodal imaging analysis of coordinated neural activation, metabolic demand, and oxygenated haemoglobin delivery in the infant brain. Confirming previous work, naturalistic social and non-social stimuli produce broad haemodynamic changes that can be refined through examining locations with coupled haemodynamic and metabolic activity (*19*). We and others have also observed broadband differences in EEG responses to social and non-social stimuli (*11*) that were also observed in the present datasets. However, examining coupling between these two phenomena uncovered a precise pattern in which a specific location at the temporal-parietal junction that differentially responds to both social and non-social stimuli was also coupled with beta and gamma band activity across chromophores in the expected pattern. We contend that this approach allows precision identification of neural specialisation through the coordination of neural, haemodynamic, and metabolic activity. Widespread use of this technique will accelerate our understanding of both the typically and atypically developing brain.

Our work is consistent with previous studies in identifying increased gamma band activity over temporal and parieto-occipital brain regions during face processing (*29*–*38*) (*39*–*42*). High-frequency neural firing is associated with localised processing (*43*) whilst lower-frequency activity is associated with larger-scale network changes and transfer of information across systems (*44*). The increase in lower-frequency activity during social attention also observed here and in other work (*11, 45*) may support larger-scale connectivity and communication of information through cross-frequency coupling (*45*). Our work further indicates that measures of metabolic load are a critical nexus in understanding localisation of brain function. Localised high-frequency activity exerts strong metabolic demand (*46, 47*) and subsequent increases in oxygenated haemoglobin (*24, 48, 49*). These increases in metabolic rate are supported by increased activity in the mitochondrial electron transport chain, resulting in the changes in cytochrome-c-oxidase we detected with broadband NIRS. Nitric oxide (which competes with oxygen to bind to cytochrome-c-oxidase) and carbon dioxide (produced as a by-product in the ETC) are key signalling molecule in controlling neurovascular coupling and thus subsequent oxygen delivery (*50, 51*). Finally, reactive oxygen species produced by the ETC are a key signal in inducing synaptic plasticity (*52*). Thus, our work is consistent with a model in which social attention induces localised high frequency brain activity in the temporal parietal junction, which increases local metabolic rates, triggering synaptic plasticity and subsequent oxygen delivery to a broader region.

Our work specifically pinpoints the importance of the temporal-parietal junction in early social brain function. Previous studies measuring haemodynamic activity have identified early sensitivity of this region to social stimuli from at least 4 months (*53*), alongside a broader network of other regions. Here, we pinpoint this specific location as having coupled neuronal, metabolic, and haemodynamic activity that is modulated in opposite directions by complex social and non-social content. In the adult brain, the temporal-parietal junction has received considerable attention and there are several competing models of its function. It has been linked to mentalising (*54, 55*) and reorienting attention to behaviourally relevant stimuli (*56*); it can be viewed as a nexus area where the convergence of attention, language, memory and social processing supports a social context for behaviour ((*57*) or as a region that is active when awareness of a prediction permits attentional control (*58*). Intriguingly, recent formulations within the predictive coding framework link the right temporal-parietal junction to a domain-general role in prediction, perhaps representing the precision of priors (*59*). Predictability has been linked to energy-efficiency, with some computational models showing that energy limitations are the only requirement for driving the emergence of predictive coding (*60*). Increases in beta/gamma have also been linked to unexpected reward processing (*61*). Taken together, our results may indicate the early presence of priors for social interaction that are being actively updated (in contrast to the dynamic toys, which may already be more predictable).

The methods we developed have extensive application in both neurotypical and atypical brain function. Assessing coupling over developmental time will indicate the mechanisms underpinning neural specialisation and constrain theoretical frameworks seeking to explain specialisation in the adult brain. The mechanisms of neurovascular coupling remain unclear in the adult brain (*50*), and are developing in infancy (*17*), and novel multimodal and non-invasive approaches to their identification could yield significant progress. Computational models could test the role of constraints in energy supply on developing localisation of function. Further, the region identified here also shows atypical haemodynamic responsiveness in infants with later symptoms of autism (*62*); since mitochondrial dysfunction has become an increasing focus in autism (*63*) the possibility that atypical coupling may impact specialisation in autism is an important hypothesis to test. Further, our methods have applicability in determining the impacts of early brain injury. Recent work (*64*) measured both cerebral oxygenation and energy metabolism in neonates with brain injury (hypoxic-ischaemic encephalopathy) and demonstrated that the relationship between metabolism and oxygenation was able to predict injury severity. This therefore provided a clinical, non-invasive biomarker of neonatal brain injury. Indicating applicability across the lifespan (*65*) simultaneous measurements of cerebral oxygenation, metabolism and neural activity in epilepsy revealed unique metabolic profiles for healthy brain regions in comparison to those with the regions of the epileptic focus. This work demonstrates the strength of combining measurements from multiple modalities to investigate brain states, particularly in clinical populations.

Our work has several limitations. We used naturalistic stimuli to maximise ecological validity; however, this reduces our ability to probe the function of the temporal-parietal junction across specific stimulus dimensions and this is an important target for future work. Limitations of current technology meant we recorded from the right hemisphere only and thus cannot determine the specificity of our findings to left temporal-parietal junction; engineering advances are required to produce whole-head bNIRS devices.

## Conclusion

Energy metabolism and neural activity are known to be tightly coupled in order to meet the high energetic demands of the brain, both during a task (*66, 67*) and at rest (*68*). It has been hypothesised that the level of correspondence between energy metabolism and neuronal activity may be an indicator for brain specialisation (*28, 66, 69*). Here, we developed a system to simultaneously measure multichannel broadband NIRS with EEG in 4-to-7-month-old infants to investigate the neurovascular and neurometabolic coupling. We presented a novel study combining bNIRS and EEG and show stimulus-dependent coupling between haemodynamic, metabolic, and neural activity in the temporal-parietal junction. The results highlight the importance of investigating the energetic basis of brain functional specialisation and opens a new avenue of research which may show high utility for studying neurodevelopmental disorders and in clinical populations where these basic mechanisms are altered.

## Acknowledgements

M.F.S was funded by the BBSRC [BB/J014567/1], the Birkbeck Institutional Strategic Support Fund (ISSF) and the ESRC (ES/V012436/1). E.J.H.J was supported by the ESRC (ES/R009368/1). E.J.H.J, M.H.J. and M.F.S. were also supported by the AIMS-2-TRIALS programmes funded by the Innovative Medicines Initiative (IMI) Joint Undertaking Grant No. 777394. This Joint Undertaking receives support from the European Union’s Horizon 2020 research and innovation programme, with in-kind contributions from the European Federation of Pharmaceutical Industries and Associations (EFPIA) companies and funding from Autism Speaks, Autistica and SFARI. I.T. was supported by the Wellcome Trust (104580/Z/14/Z). S.L.F was supported by a UKRI Future Leaders Fellowship (MR/S018425/1) and S.L.F and C.E.E received support from the Bill and Melinda Gates Foundation (OPP1127625). M.H.J received support from the UK Medical Research Council (MR/K021389/1 & MR/T003057/1). S.B. was supported by the Progetto STARS Grants 2017 (C96C18001930005) from the University of Padova.

The work presented herein was conducted at the Centre for Brain and Cognitive Development, Birkbeck College, University of London. We are grateful to all the families who participated in this research and all the undergraduate students who assisted with data collection.

## Author Contributions

M.F.S. conducted the study. M.F.S., S.L.F., E.J.H.J, C.E.E. and M.H.J. developed the protocols for the study. I.T. provided the NIRS system and support with data acquisition.

M.F.S and S.B. analysed the data with support from P.P., S.L.F., I.T., E.J.H.J. C.E.E. and M.H.J. M.F.S. and E.J.H.J. wrote the manuscript with support from P.P., I.T., S.L.F. and M.H.J.

## Declaration of Interests

The authors declare that the research was conducted in the absence of any commercial or financial relationships that could be construed as a potential conflict of interest.

## Data availability statement

The data contains human subject data from minors and guardians provided informed consent to having data shared only with researchers involved in the project, in anonymised form. A Patient and Public Involvement (PPI) initiative at the Centre for Brain and Cognitive Development aimed to actively work in partnership with parents and guardians participating in research studies to help design and manage future research. A comprehensive public survey was conducted as part of this initiative which aimed to evaluate parent attitudes to data sharing in developmental science. This survey revealed that majority of parents do not want their data to be shared openly but are open to the data being shared with other researchers related to the project. Therefore, in order to adhere to participant preference/choice, a curated data sharing approach must be followed wherein the data can only be made available upon reasonable request through a formal data sharing and project affiliation agreement. The researcher will have to contact MFS and complete a project affiliation form providing their study aims, a detailed study proposal, plan for the analysis protocol, ethics, and plans for data storage and protection. Successful proposals will have aims aligned with the aims of the original study. Raw NIRS data, EEG data and integrated NIRS-EEG data can be made available in anonymised form. ID numbers linking the NIRS and EEG data, however, cannot be provided as parents/guardians have consented only to data being shared in anonymised form. All code used to analyse the NIRS data and the integration of the NIRS and EEG data is available on GitHub (https://github.com/maheensiddiqui91/NIRS-EEG). EEG data was processed using EEGlab which is a publicly available toolbox.

## Methods

### Participants

The study protocol was approved by the Birkbeck Ethics Committee. Participants were forty-two 4-to-7-month-old infants (mean age: 179± 16 days; 22 males and 20 females); parents provided written informed consent to participate in the study, for the publication of the research and additionally for the publication and use of any photographs taken during the study of the infant wearing the NIRS-EEG headgear. Inclusion criteria included term birth (37 – 40 weeks); exclusion criteria included known presence or family history of developmental disorders. The sample size was determined by performing a power analysis of existing data using G*Power.

### Experimental Procedure

The experimental stimuli were designed using Psychtoolbox in Matlab (Mathworks, USA) and consisted of social and non-social videos. The social videos consisted of a variety of full-colour video clips of actors performing nursery rhymes such as “pat-a-cake” and “wheels on the bus”. The non-social videos consisted of dynamic video clips of moving mechanical toys. The visual and auditory components of both social and non-social videos was matched. These videos have been used extensively in prior infant studies in both EEG studies (*11*) and NIRS studies (*70, 71*). Both social and non-social experimental conditions were presented alternatingly for a varying duration between 8-12 s. The baseline condition consisted of static transport images, for example cars and helicopters, which were presented for a pseudorandom duration of 1 – 3 s each for a total of 8 s. Following the presentation of the baseline condition, a fixation cross in the shape of a ball or a flower appeared in the centre of the screen to draw the infant’s attention back to the screen in case the infant had become bored during the baseline period. The following experimental condition was then presented once the infant’s attention was on the fixation cross. **Error! Reference source not found**.a depicts the order of stimulus presentation. All infants sat in their parent’s lap at an approximate distance of 65 cm from a 35-in screen which was used to display the experimental stimuli. The study began with a minimum 10 s rest period to draw the infant’s attention towards the screen during which the infant was presented with various shapes in the four corners of the screen. Following this, the baseline and experimental stimuli were presented alternatingly until the infant became bored or fussy.

### Data acquisition and array placement

bNIRS and EEG data was acquired simultaneously and the bNIRS optodes and EEG electrodes were positioned on the head using custom-built, 3-D printed arrays which were embedded within a soft neoprene cap (Neuroelectrics, Spain). Figures 5a and 5b show the locations of bNIRS optodes and EEG electrodes on the head. Figure 1b shows the combined bNIRS-EEG headgear positioned on an infant. The array was designed to allow measurement from several cortical regions which included occipital, parietal, temporal, and central regions to allow investigation of neurovascular coupling in different cortical regions that are expected to be activated by dynamic stimuli.

**Figure 5:**
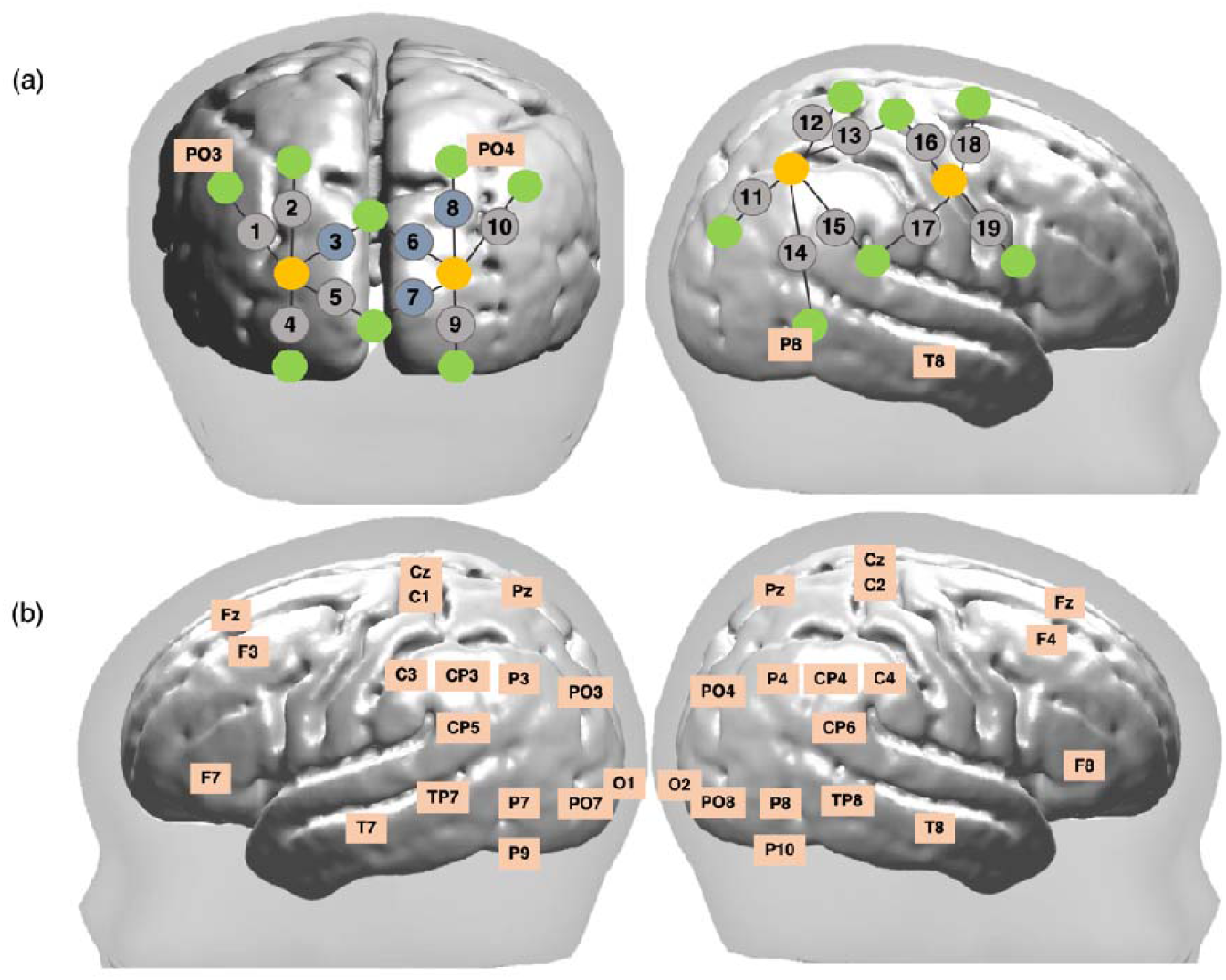
Schematic representation of bNIRS and EEG channel locations. (a) Locations of bNIRS channels (grey circles) over the occipital cortex and the right hemisphere and locations of the bNIRS sources (orange circles) and detectors (green circles) relative to EEG 10/20 locations. Channels shown in blue (3, 6, 8 and 10) were excluded from the analysis (b) Locations of the 32 EEG electrodes.

### Broadband NIRS

Brain haemodynamic (Δ[HbO_2_], Δ[HHb]) and metabolic changes (Δ[oxCCO]) were measured using an in-house broadband NIRS system developed at University College London (*72*). The bNIRS system consisted of two light sources that consisting of halogen light bulbs (Phillips) that emitted light in the near-infrared range (504 – 1068 nm). The light was directed to the infant’s head through customised bifurcated optical fibres (Loptek, Germany), allowing each light source to split into two pairs of light sources. This formed a total of four light sources at the participant-end and each pair of light sources were controlled by a time multiplexing mechanism whereby one pair of light sources was on every 1.4 s. The system also consisted of fourteen detector fibres at the participant-end which were connected to two spectrometers, seven for each spectrometer (in-house developed lens spectrographs and PIXIS512f CCD cameras (Princeton Instruments). The configuration of four light sources and fourteen detectors formed a total of nineteen measurement channels. These were positioned over the occipital cortex and the right hemisphere as shown in Figure 5a. The source-detector separation was 2.5 cm.

Data were analysed in Matlab (Mathworks, USA) using in-house scripts. First, for each participant, across all wavelengths, wavelet-based motion correction (*73*) was applied to the attenuation change signal to correct for motion artifacts. The tuning parameter α□=□0.8 was used. Following this, the UCLn algorithm (*18*) was used with a wavelength-dependent, age-appropriate fixed differential path-length factor (DPF) value of 5.13 (*74*). Changes in concentration of HbO_2_, HHb and oxCCO were calculated using 120 wavelengths between 780 – 900 nm. A 4^th^-order bandpass Butterworth filter from 0.01 – 0.4 Hz was used to filter the data. For each infant, channels were assessed for signal quality and any channels with poor signal quality were rejected. Following this, the HbO_2_, HHb and oxCCO time-series were entered into a General Linear Model (GLM) to correlate bNIRS and EEG data.

For each infant, intensity counts (or photon counts) from each of the fourteen detectors were used to assess the signal-to-noise (SNR) ratio at each channel and the channels with intensity counts lower than 2000 or higher than 40,000 were excluded (*72*). If an infant had more than 60% of channels excluded, they were excluded from the study. At the group level, five channels over the occipital cortex were excluded due to poor SNR in majority of infants (Channel 3 excluded in 64% of infants, Channel 6 excluded in 83% of infants, Channel 7 excluded in 64% of infants, Channel 8 excluded in 79% of infants) and one channel over the right hemisphere was excluded in 100% of infants due to a damaged optical fibre.

### EEG

EEG was used to measure neural activity simultaneously to haemodynamic and metabolic activity using the Enobio EEG system (Neuroelectrics, Spain) which is a wireless gel-based system. The system consisted of 32 electrodes, the locations of which are shown in Figure 5b. The sampling rate of the system was 500 Hz. The experimental protocol in Psychtoolbox sent event markers to both bNIRS and EEG systems using serial port communication which was then used to synchronise the bNIRS and EEG.

All data were analysed using the EEGlab Toolbox (Schwartz Centre for Computation Neuroscience, UC San Diego, USA) and in-house scripts in Matlab (Mathworks, USA). The raw EEG signal was band-pass filtered between 0.1 – 100 Hz and a notch filter (48 – 52 Hz) was applied to remove artifacts due to line noise. Following this, blocks of the data were created such that they consisted of the baseline period prior to the stimulus presentation and the entire following stimulus period. These blocks were then segmented into 1 s segments such that for both the baseline and the stimulus, each 8 – 12 s presentation of the baseline condition or the stimulus condition yielded 8 – 12 × 1 s segments. These 1 s segments consisted of 200 ms of the previous 1 s segment and 800 ms of the current segment and the 200 ms was used for baseline correction of each 1 s segment. Segments where the infants were not visually attending to the stimulus were removed. Artifacts were detected using automatic artifact-detection in EEGlab and through manual identification. EEG segments were rejected if the signal amplitude exceeded 200 μV, or if electro-ocular, movement, or muscular artifacts occurred. Channels with noisy data were interpolated by an algorithm incorporated within EEGlab. Data were then re-referenced to the average reference.

Within each block (consisting of the baseline period and the stimulus period), each artifact-free 1 s segment was subjected to a power analysis to calculate the average root mean square (RMS) power for both low and high frequency bands – theta (3 – 6 Hz), alpha (8 – 12 Hz), beta (13 – 30 Hz) and gamma (20 – 60 Hz), within each 1 s segment. This then yielded the average RMS power across the block (baseline period + following stimulus period). Baseline correction was performed by subtracting the average of the 2 s of the baseline period from the entire block. RMS power was chosen as the metric to correlate bNIRS and EEG data as previous studies have demonstrated that task-related BOLD changes are best explained by RMS (*75, 76*). The blocks were then averaged across trials to obtain an averaged RMS response per participant. A portion of the averaged RMS power was then entered into a GLM analysis described below – this consisted of two seconds of the baseline period and 8 seconds of the stimulus period. Figure 6a provides a visual depiction of how the RMS power was derived from the pre-processed EEG data. For each participant, the RMS power was also averaged across the stimulus period for statistical analysis of the EEG data. For each frequency band, statistical t-tests were performed on this averaged RMS power comparing the social condition versus the baseline (RMS power was averaged during the baseline period), the non-social condition versus the baseline and social versus non-social. The false discovery rate (FDR) procedure using the Benjamin Hochberg method (*77*) was performed to correct for multiple comparisons.

**Figure 6:**
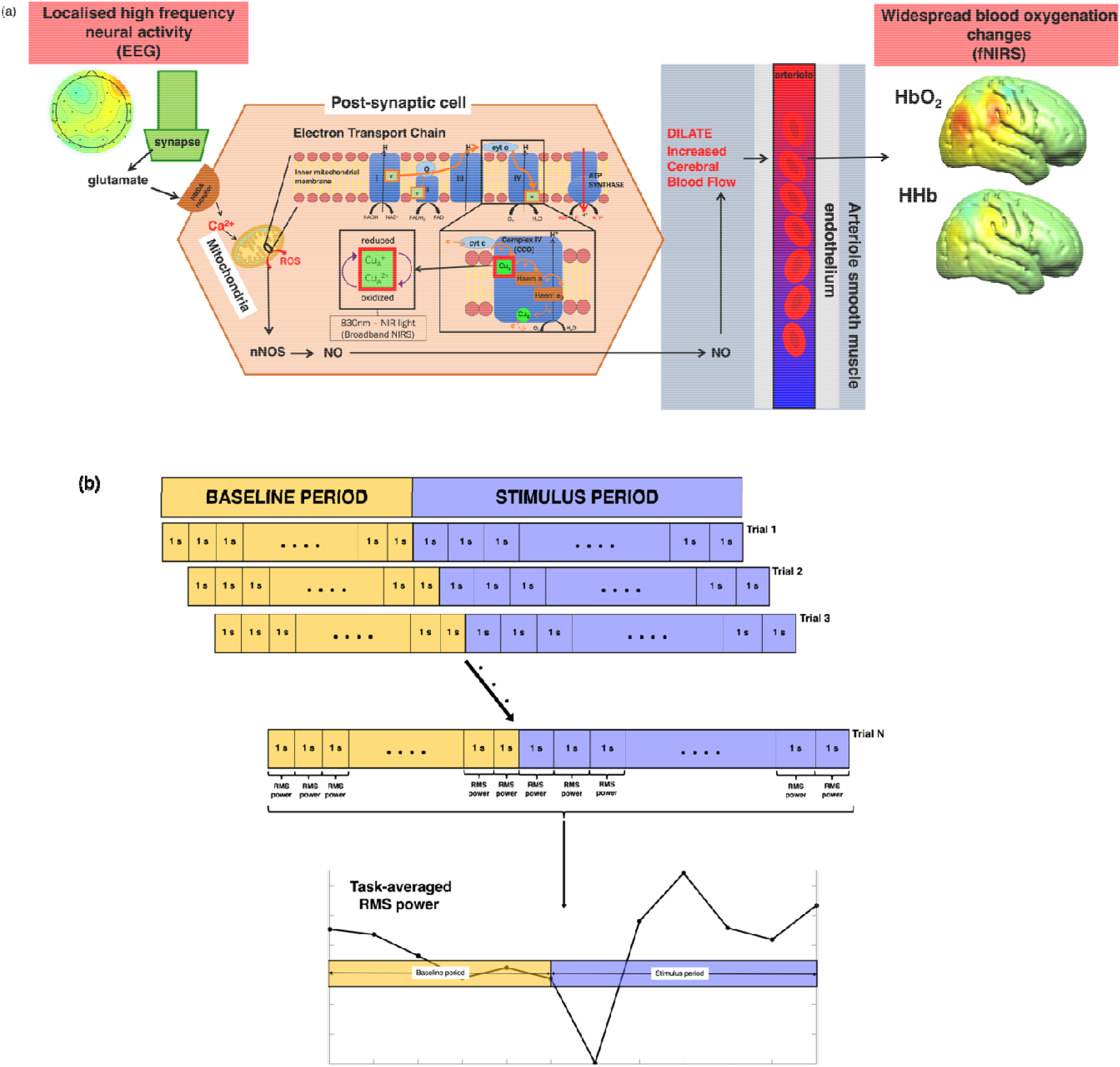
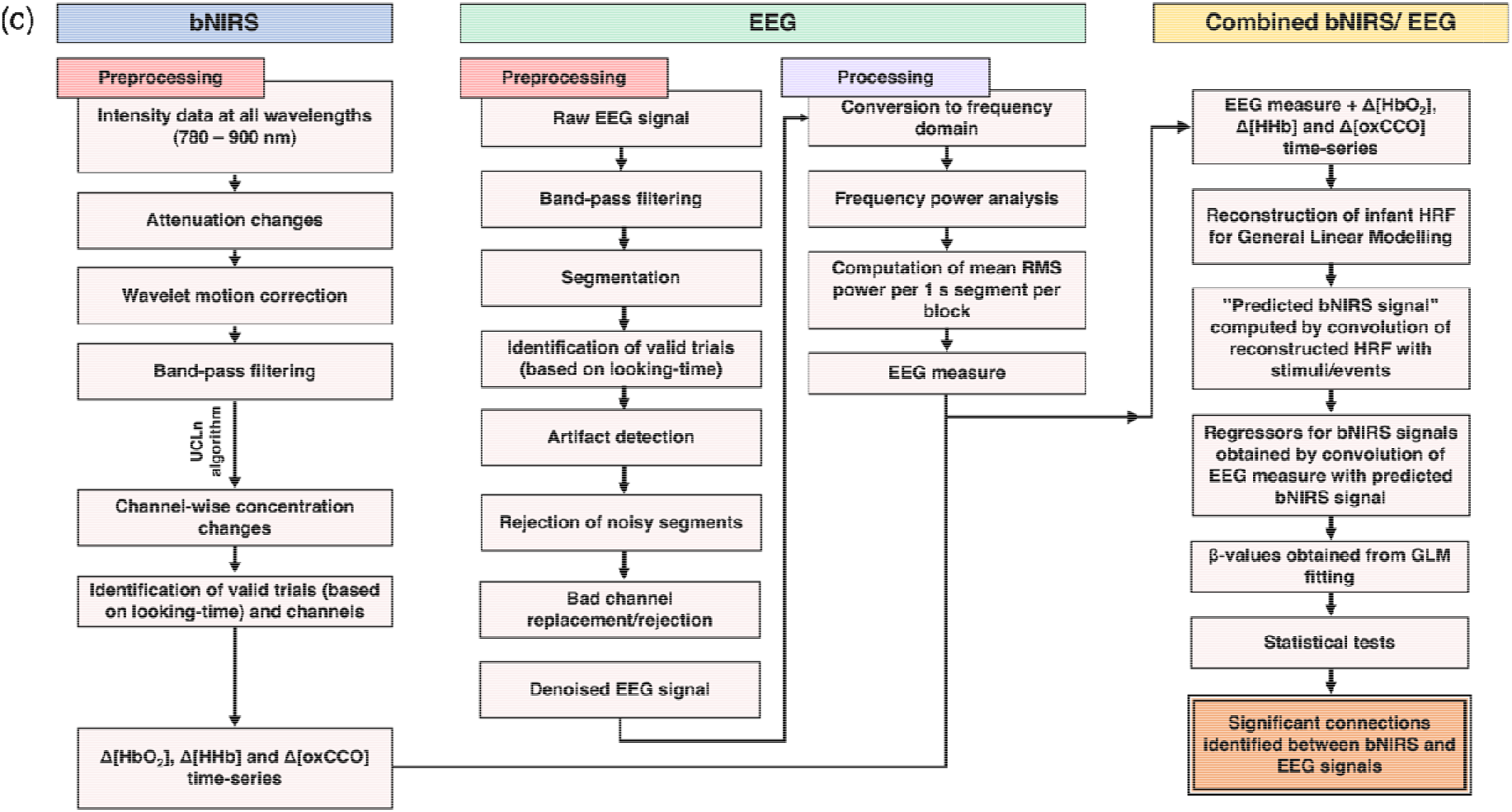
(a) Summary of the signalling pathways that mediate neurovascular coupling. High-frequency neural activity causes the release the excitatory neurotransmitter glutamate which binds to N-methyl-D-aspartate (NMDA) receptors in interneurons. This causes an influx of calcium (Ca^2+^) which in turn leads to an increase in ATP production through the mitochondrial electron transport chain (ETC). As a by-product, nitric oxide (NO) and reactive oxygen species (ROS) are produced. NO dilates arterioles to increase blood flow leading to increased oxygen delivery in surrounding brain regions. ROS influence synaptic plasticity. (b) Procedure for deriving the EEG RMS power from the pre-processed EEG data. The task-averaged RMS power shown here is average theta power across all infants from a single channel for the purposes of outlining the procedure (c) Flow chart for the data analysis pipelines for bNIRS (left), EEG (middle) and combined bNIRS-EEG (right).

### Data Analysis

Figure 6b outlines the data analysis pipelines for both bNIRS and EEG data, as well as the procedure for the combined bNIRS-EEG analysis.

### Combined NIRS-EEG analysis

A GLM (*78*) approach was employed to investigate the relationship between the bNIRS hemodynamic and metabolic data with the EEG neural data. The canonical GLM typically uses a model of the expected haemodynamic response, i.e. the hemodynamic response function (HRF), to predict the hemodynamic signal. However, given the differences in the haemodynamic response in adults and infants, the standard adult HRF model cannot be assumed for infant data. For example, infants display a delay in their haemodynamic responses (*79*–*81*). In addition, the analogous of the HRF is not established for the metabolic response (i.e. the metabolic response function or MRF). Therefore, the first step of this analysis involved reconstructing the HRF for HbO_2_ and HHb and the MRF for oxCCO before combing bNIRS and EEG data.

The reconstruction of the infant HRF and MRF started with block-averaging the HbO_2_, HHb, and oxCCO signals for social and non-social conditions within each baby. Based on our previous study (*19*), we selected only the channels that displayed statistically significant responses to the contrast task versus baseline. The single subjects block-averaged responses were averaged across the social and non-social conditions and then across the significant channels. The resulting block-averaged responses were then averaged across the group to obtain a “grand average” HbO_2_, HHb and oxCCO response.

The grand average was then used in an iterative approach to estimate the HRF and MRF that best fit the HbO_2_, HHb and oxCCO responses. This involved fitting the grand averaged signals with different HRF/MRF models starting from the canonical HRF made of two gamma functions and varying the following parameters: 1) delay of response, 2) delay of the undershoot and 3) ratio of response to undershoot to identify the combination of parameters that best reconstructed the infant HRF/MRF for the social/non-social stimuli. The parameters were varied in increments of 1 s such that the delay of the response was varied from 5 s to 15 s from the stimulus onset, the delay of the undershoot was varied from 5 to 20 s and the ratio of the response to the undershoot was varied from 2 to 6 s. All possible combinations of parameters were tested. The grand average responses were fitted with each HRF/MRF in GLM approach, and β-values were obtained for each combination of the HRF/MRF parameters. The β-values were entered into a statistical test and the parameter combinations that yielded the highest, statistically significant β-values (i.e. the model best fitting the data) were selected to reconstruct the infant HRF/MRF. This is approach is similar to those used previously to reconstruct the infant HRF (*81*) and identified the best fit to be with a 2-s delay of response for HbO_2_ and HHb and a 3-s delay of response for oxCCO in comparison to the adult HRF (i.e. 6 s). Moreover, the delay of the undershoot was 9-s earlier for all chromophores and the ratio of the response to the undershoot was 2 for HbO_2_ and HHb and 3 for oxCCO, in comparison to 6 for the adult HRF. The new reconstructed HRF was then used for the GLM approach to correlate bNIRS and EEG data. The process for estimating the HRF and MRF has been depicted in Figure 7.

**Figure 7:**
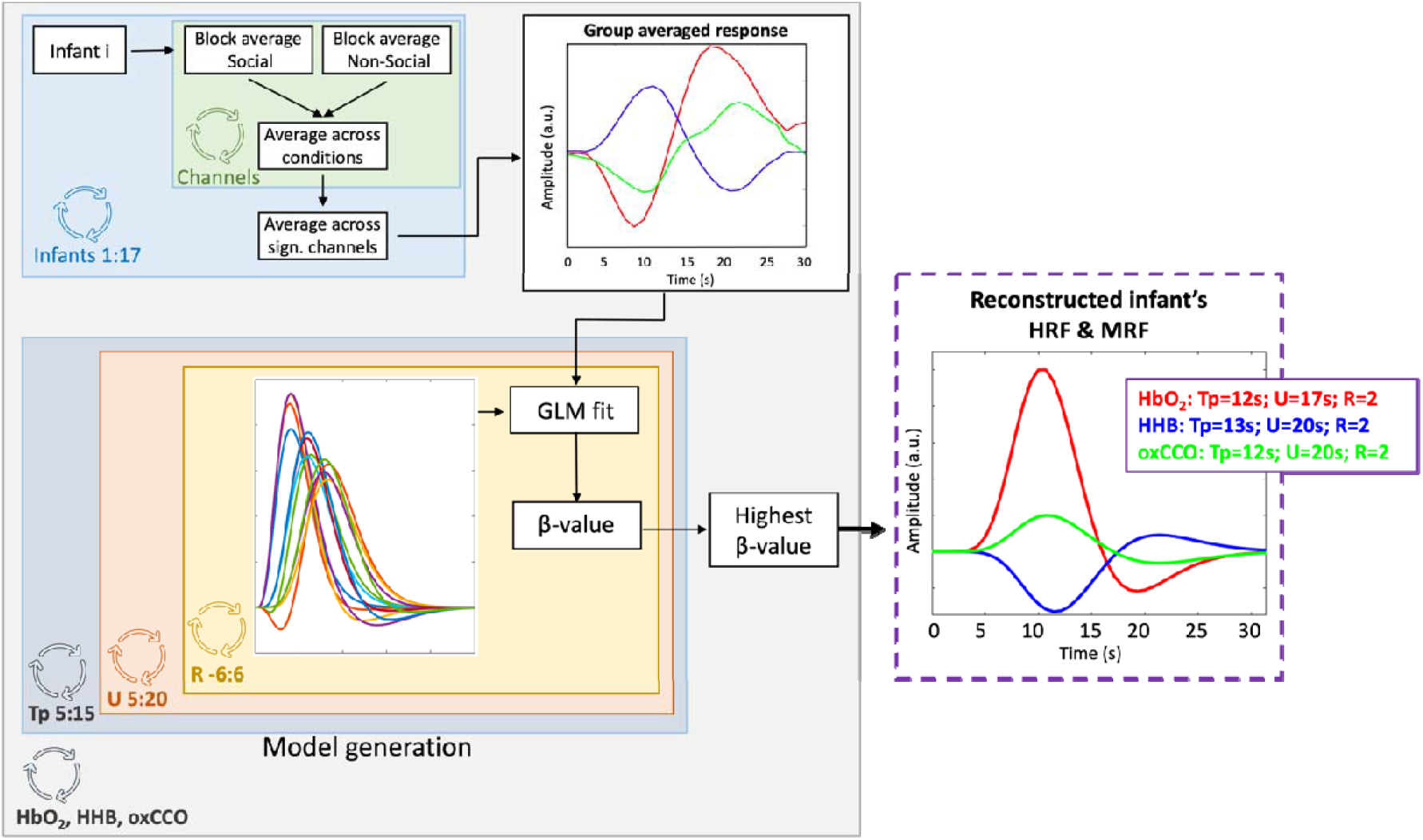
Procedure for obtaining the reconstructed haemodynamic response function (HRF) and the metabolic response function (MRF).

To constrain the analysis, we chose to investigate coupling of haemodynamic and metabolic with neural activity at specific channels. For this, we used the results from an analysis we described previously that combined bNIRS haemodynamic and metabolic signals (*19, 27*). The results from this identified task-relevant cortical regions that displayed high levels of haemodynamic and metabolic coupling. The bNIRS channels that displayed significant haemodynamic and metabolic coupling for social and non-social conditions were used here. All EEG channels were used as EEG is not as spatially specific as bNIRS. For each infant, for each chromophore, for each channel and each EEG frequency band, the new infant HRF/MRF that was reconstructed in the previous step was convolved with the events to obtain the “predicted” bNIRS signal. The “predicted” bNIRS signal was then convolved with the EEG RMS power block (consisting only of the data from the stimulus period) at each frequency band to obtain the neural regressor for the bNIRS data, considering both social and non-social conditions together. The design matrix thus included the neural regressor reflecting the increased in EEG activity to the social and non-social stimuli and used to fit the bNIRS data. This was performed for HbO2, HHb, and oxCCO individually for all the channels. β-values were estimated for each channel and t-tests against 0 were conducted to test whether there was a statistically significant association between bNIRS signals and EEG frequency bands. The false discovery rate (FDR) procedure using the Benjamin Hochberg method (*77*) was performed to correct for multiple comparisons. The FDR-corrected significant t-values were plotted. This method has been used in numerous studies previously in correlating fMRI BOLD – EEG (*20*). For each frequency band, FDR-corrected, significant, β-values were also averaged (1) for bNIRS and EEG channels over the right hemisphere and (2) between bNIRS channels in the right hemisphere and EEG channels in the left hemisphere to obtain an estimate of the frequency band where bNIRS and EEG activity associated most strongly within hemispheres and across hemispheres. Only bNIRS channels that displayed significant (prior to FDR correction) haemodynamic and metabolic coupling were used for this analysis (as indicated in Figure and **Error! Reference source not found**.). For the social condition, these were channels 4, 12, 13 and 14 for HbO_2_, channels 11, 12, 14 and 18 for HHb and channels 4, 11, 12, 13, 14 and 18 for oxCCO. For the non-social condition, these were channels 5, 12 and 14 for HbO_2_, channels 5, 12, 14 and 16 for HHb and channels 5, 12, 14 and 16 for oxCCO.

For the bNIRS analysis, data from 25 infants was included while for the EEG analysis, data from 35 infants were included. For the joint bNIRS-EEG analysis, only infants that had both valid bNIRS and EEG data for social and non-social conditions were included and therefore 17 infants were included in this analysis.

### Image reconstruction

Image reconstruction was performed on the bNIRS data, at the individual subject level and then averaged across infants to produce a grand average that is shown in Figure 4. For this analysis, three additional long-distance channels were created over the right hemisphere (in addition to the 19 bNIRS channels) that had a source/detector separation of 4.3cm.

For this analysis, the block averaged attenuation changes at 13 discrete wavelengths (from 780 – 900 nm at 10 nm intervals) for each infant were selected from the bNIRS data. This was done to reduce the computational burden of the reconstruction (*82*). A four-layer infant head-model (consisting of the grey matter (GM), white matter (WM), cerebrospinal fluid (CSF) and extra cerebral tissue) was built using averaged MRI data from a cohort of 12-month-old infants presented in Shi et al. (*83*). The Betsurf segmentation procedure (*84*) was then used to define an outer scalp boundary from the average head MRI template. The voxelised four-layer model was converted to a high-resolution tetrahedral mesh (∼7.8 × 10^5^ nodes and ∼4.7 × 10^6^ elements) using the iso2mesh software (Fang & Boas, 2009). The same software was used to create the GM surface mesh (∼5.8 × 10^4^ nodes and ∼1.2 × 10^5^ faces), used to visualise the reconstructed images.

The reconstruction of images of HbO_2_, HHb and ΔoxCCO are described elsewhere (*85*), using a multispectral approach (*86*). Wavelength-specific Jacobians were computed with the Toast++ software (*87*) on the tetrahedral head mesh and projected onto a 50 × 60 × 50 voxel regular grid for reconstruction, using an intermediate finer grid of 100 × 120 × 100 voxels to optimize the mapping between mesh and voxel space. Optical properties were assigned to each tissue type and for each wavelength by fitting all published values for these tissue types (*88*–*90*). Diffuse boundary sources and detectors were simulated as a Gaussian profile with a 2-mm standard deviation, and Neumann boundary conditions were applied. The inverse problem was solved employing the LSQR method to solve the matrix equations resulting from the minimization and using first-order Tikhonov regularization, with the parameter covariance matrix containing the diagonal square matrices with the background concentration values of the three chromophores (23.7 for HbO_2_, 16 for HHb and 6 for ΔoxCCO) (*91, 92*) and the noise covariance matrix set as the identity matrix. The maximum number of iterations allowed to the LSQR method was set to 50, and with a tolerance of 10^−5^. The regularization hyperparameter λ was set to 10^−2^.

The reconstructed images, defined on the same regular grid of the Jacobian, were remapped to the tetrahedral head mesh and then projected to the GM surface mesh, by assigning a value to each node on the GM boundary surface that was equal to the mean value of all the tetrahedral mesh node values within a 3-mm radius. The concentration changes for HbO_2_ and HHb were normalised to the maximum concentration change of HbO_2_ while ΔoxCCO was normalised to its own maximum change in concentration.

## Notes

### Competing Interest Statement

The authors have declared no competing interest.

## References

1. M. H. Johnson, Functional brain development in humans. Nat. Rev. Neurosci. 2, 475– 483 (2001).

2. C. C. Goren, M. Sarty, P. Y. K. Wu, Visual following and pattern discrimination of face like stimuli by newborn infants. Pediatrics. 56, 544–549 (1975).

3. M. H. Johnson, Interactive specialization: a domain-general framework for human functional brain development? Dev. Cogn. Neurosci. 1, 7–21 (2011).

4. N. Uomini, J. Fairlie, R. D. Gray, M. Griesser, Extended parenting and the evolution of cognition. Philos. Trans. R. Soc. B Biol. Sci. 375, 20190495 (2020).

5. D. P. Kennedy, R. Adolphs, The social brain in psychiatric and neurological disorders. Trends Cogn. Sci. 16, 559–572 (2012).

6. M. Eimer, The Face-Sensitivity of the N170 Component. Front. Hum. Neurosci. 5 (2011), doi:10.3389/fnhum.2011.00119.

7. S. Conte, J. E. Richards, M. W. Guy, W. Xie, J. E. Roberts, Face-sensitive brain responses in the first year of life. NeuroImage. 211, 116602 (2020).

8. M. de Haan, M. H. Johnson, H. Halit, Development of face-sensitive event-related potentials during infancy: a review. Int. J. Psychophysiol. Off. J. Int. Organ. Psychophysiol. 51, 45–58 (2003).

9. H. L. Kosakowski, M. A. Cohen, A. Takahashi, B. Keil, N. Kanwisher, R. Saxe, Selective responses to faces, scenes, and bodies in the ventral visual pathway of infants. Curr. Biol. 32, 265-274.e5 (2022).

10. T. Farroni, A. M. Chiarelli, S. Lloyd-Fox, S. Massaccesi, A. Merla, V. Di Gangi, T. Mattarello, D. Faraguna, M. H. Johnson, Infant cortex responds to other humans from shortly after birth. Sci. Rep. 3 (2013), doi:10.1038/srep02851.

11. E. J. H. Jones, K. Venema, R. Lowy, R. K. Earl, S. J. Webb, Developmental changes in infant brain activity during naturalistic social experiences. Dev. Psychobiol. 57, 842– 853 (2015).

12. M. E. Raichle, M. A. Mintun, Brain work and brain imaging. Annu. Rev. Neurosci. 29, 449–476 (2006).

13. L. Sokoloff, Energetics of functional activation in neural tissues. Neurochem. Res. 24, 321–329 (1999).

14. P. Steiner, Brain Fuel Utilization in the Developing Brain. Ann. Nutr. Metab. 75, 8–18 (2019).

15. R. Leech, G. Scott, R. Carhart-Harris, F. Turkheimer, S. D. Taylor-Robinson, D. J. Sharp, Spatial Dependencies between Large-Scale Brain Networks. PLOS ONE. 9, e98500 (2014).

16. S. Vaynman, Z. Ying, A. Wu, F. Gomez-Pinilla, Coupling energy metabolism with a mechanism to support brain-derived neurotrophic factor-mediated synaptic plasticity. Neuroscience. 139, 1221–1234 (2006).

17. M. Kozberg, E. Hillman, Neurovascular coupling and energy metabolism in the developing brain. Prog. Brain Res. 225, 213–242 (2016).

18. G. Bale, C. E. Elwell, I. Tachtsidis, From Jöbsis to the present day: a review of clinical near-infrared spectroscopy measurements of cerebral cytochrome-c-oxidase. J. Biomed. Opt. 21, 91307 (2016).

19. M. F. Siddiqui, P. Pinti, S. Lloyd-Fox, E. J. H. Jones, S. Brigadoi, L. Collins-Jones, I. Tachtsidis, M. H. Johnson, C. E. Elwell, Regional Haemodynamic and Metabolic Coupling in Infants. Front. Hum. Neurosci. 15 (2022), doi:10.3389/fnhum.2021.780076.

20. R. Scheeringa, P. Fries, K.-M. Petersson, R. Oostenveld, I. Grothe, D. G. Norris, P. Hagoort, M. C. M. Bastiaansen, Neuronal Dynamics Underlying High- and Low-Frequency EEG Oscillations Contribute Independently to the Human BOLD Signal. Neuron. 69, 572–583 (2011).

21. R. Scheeringa, K. M. Petersson, R. Oostenveld, D. G. Norris, P. Hagoort, M. C. M. Bastiaansen, Trial-by-trial coupling between EEG and BOLD identifies networks related to alpha and theta EEG power increases during working memory maintenance. NeuroImage. 44, 1224–1238 (2009).

22. Stern, Simultaneous EEG and fMRI of the alpha rhythm. NeuroReport, 2487–2492 (2002).

23. H. Yuan, T. Liu, R. Szarkowski, C. Rios, J. Ashe, B. He, Negative covariation between task-related responses in alpha/beta-band activity and BOLD in human sensorimotor cortex: An EEG and fMRI study of motor imagery and movements. NeuroImage. 49, 2596–2606 (2010).

24. J. Niessing, Hemodynamic Signals Correlate Tightly with Synchronized Gamma Oscillations. Science. 309, 948–951 (2005).

25. N. K. Logothetis, J. Pauls, M. Augath, T. Trinath, A. Oeltermann, Neurophysiological investigation of the basis of the fMRI signal. Nature. 412, 150–157 (2001).

26. S. P. Koch, P. Werner, J. Steinbrink, P. Fries, H. Obrig, J. Neurosci., in press, doi:10.1523/JNEUROSCI.1402-09.2009.

27. P. Pinti, M. F. Siddiqui, A. D. Levy, E. J. H. Jones, I. Tachtsidis, An analysis framework for the integration of broadband NIRS and EEG to assess neurovascular and neurometabolic coupling. Sci. Rep. 11, 3977 (2021).

28. E. Shokri-Kojori, D. Tomasi, B. Alipanahi, C. E. Wiers, G.-J. Wang, N. D. Volkow, Correspondence between cerebral glucose metabolism and BOLD reveals relative power and cost in human brain. Nat. Commun. 10, 690 (2019).

29. S. Uono, W. Sato, T. Kochiyama, Y. Kubota, R. Sawada, S. Yoshimura, M. Toichi, Time course of gamma-band oscillation associated with face processing in the inferior occipital gyrus and fusiform gyrus: A combined fMRI and MEG study. Hum. Brain Mapp. 38, 2067–2079 (2017).

30. A. S. Ghuman, N. M. Brunet, Y. Li, R. O. Konecky, J. A. Pyles, S. A. Walls, V. Destefino, W. Wang, R. M. Richardson, Dynamic encoding of face information in the human fusiform gyrus. Nat. Commun. 5, 5672 (2014).

31. A. D. Engell, G. McCarthy, The Relationship of Gamma Oscillations and Face-Specific ERPs Recorded Subdurally from Occipitotemporal Cortex. Cereb. Cortex. 21, 1213– 1221 (2011).

32. F. Bossi, I. Premoli, S. Pizzamiglio, S. Balaban, P. Ricciardelli, D. Rivolta, Theta- and Gamma-Band Activity Discriminates Face, Body and Object Perception. Front. Hum. Neurosci. 14, 74 (2020).

33. D. Anaki, E. Zion-Golumbic, S. Bentin, Electrophysiological neural mechanisms for detection, configural analysis and recognition of faces. NeuroImage. 37, 1407–1416 (2007).

34. A. Ishai, L. Ungerleider, A. Martin, J. Haxby, The Representation of Objects in the Human Occipital and Temporal Cortex. J. Cogn. Neurosci. 12 Suppl 2, 35–51 (2000).

35. K. A. Pelphrey, J. P. Morris, G. McCarthy, Neural basis of eye gaze processing deficits in autism. Brain. 128, 1038–1048 (2005).

36. W. Sato, T. Kochiyama, S. Uono, K. Matsuda, K. Usui, Y. Inoue, M. Toichi, Rapid, high-frequency, and theta-coupled gamma oscillations in the inferior occipital gyrus during face processing. Cortex. 60, 52–68 (2014).

37. E. Zion-Golumbic, T. Golan, D. Anaki, S. Bentin, Human face preference in gamma-frequency EEG activity. NeuroImage. 39, 1980–1987 (2008).

38. Z. Gao, A. Goldstein, Y. Harpaz, M. Hansel, E. Zion-Golumbic, S. Bentin, A magnetoencephalographic study of face processing: M170, gamma-band oscillations and source localization. Hum. Brain Mapp. 34, 1783–1795 (2013).

39. M. Bayer, M. T. Rubens, T. Johnstone, Simultaneous EEG-fMRI reveals attention-dependent coupling of early face processing with a distributed cortical network. Biol. Psychol. 132, 133–142 (2018).

40. M. Müller-Bardorff, M. Bruchmann, M. Mothes-Lasch, P. Zwitserlood, I. Schlossmacher, D. Hofmann, W. Miltner, T. Straube, Early brain responses to affective faces: A simultaneous EEG-fMRI study. NeuroImage. 178, 660–667 (2018).

41. V. T. Nguyen, R. Cunnington, The superior temporal sulcus and the N170 during face processing: Single trial analysis of concurrent EEG–fMRI. NeuroImage. 86, 492–502 (2014).

42. V. T. Nguyen, M. Breakspear, R. Cunnington, Fusing concurrent EEG–fMRI with dynamic causal modeling: Application to effective connectivity during face perception. NeuroImage. 102, 60–70 (2014).

43. A. von Stein, J. Sarnthein, Different frequencies for different scales of cortical integration: from local gamma to long range alpha/theta synchronization. Int. J. Psychophysiol. 38, 301–313 (2000).

44. R. T. Canolty, R. T. Knight, The functional role of cross-frequency coupling. Trends Cogn. Sci. 14, 506–515 (2010).

45. B. van der Velde, T. White, C. Kemner, The emergence of a theta social brain network during infancy. NeuroImage. 240, 118298 (2021).

46. A. J. Smith, H. Blumenfeld, K. L. Behar, D. L. Rothman, R. G. Shulman, F. Hyder, Cerebral energetics and spiking frequency: The neurophysiological basis of fMRI. Proc. Natl. Acad. Sci. U. S. A. 99, 10765–10770 (2002).

47. O. Kann, The Energy Demand of Fast Neuronal Network Oscillations: Insights from Brain Slice Preparations. Front. Pharmacol. 2 (2012) (available at https://www.frontiersin.org/articles/10.3389/fphar.2011.00090).

48. N. K. Logothetis, J. Pauls, M. Augath, T. Trinath, A. Oeltermann, Neurophysiological investigation of the basis of the fMRI signal. Nature. 412, 150–157 (2001).

49. J. B. M. Goense, N. K. Logothetis, Neurophysiology of the BOLD fMRI Signal in Awake Monkeys. Curr. Biol. 18, 631–640 (2008).

50. P. S. Hosford, A. V. Gourine, What is the key mediator of the neurovascular coupling response? Neurosci. Biobehav. Rev. 96, 174–181 (2019).

51. P. S. Hosford, J. A. Wells, S. Nizari, I. N. Christie, S. M. Theparambil, P. A. Castro, A. Hadjihambi, L. F. Barros, I. Ruminot, M. F. Lythgoe, A. V. Gourine, CO2 signaling mediates neurovascular coupling in the cerebral cortex. Nat. Commun. 13, 2125 (2022).

52. M. C. Oswald, P. S. Brooks, M. F. Zwart, A. Mukherjee, R. J. West, C. N. Giachello, K. Morarach, R. A. Baines, S. T. Sweeney, M. Landgraf, Reactive oxygen species regulate activity-dependent neuronal plasticity in Drosophila. eLife. 7, e39393 (2018).

53. S. Lloyd-Fox, K. Begus, D. Halliday, L. Pirazzoli, A. Blasi, M. Papademetriou, M. K. Darboe, A. M. Prentice, M. H. Johnson, S. E. Moore, C. E. Elwell, Cortical specialisation to social stimuli from the first days to the second year of life: A rural Gambian cohort. Dev. Cogn. Neurosci. 25, 92–104 (2017).

54. M. Schurz, J. Radua, M. Aichhorn, F. Richlan, J. Perner, Fractionating theory of mind: A meta-analysis of functional brain imaging studies. Neurosci. Biobehav. Rev. 42, 9–34 (2014).

55. M. Schurz, M. G. Tholen, J. Perner, R. B. Mars, J. Sallet, Specifying the brain anatomy underlying temporo-parietal junction activations for theory of mind: A review using probabilistic atlases from different imaging modalities. Hum. Brain Mapp. 38, 4788– 4805 (2017).

56. M. Corbetta, G. L. Shulman, Control of goal-directed and stimulus-driven attention in the brain. Nat. Rev. Neurosci. 3, 201–215 (2002).

57. R. M. Carter, S. A. Huettel, A nexus model of the temporal-parietal junction. Trends Cogn. Sci. 17, 328–336 (2013).

58. A. I. Wilterson, S. A. Nastase, B. J. Bio, A. Guterstam, M. S. A. Graziano, Attention, awareness, and the right temporoparietal junction. Proc. Natl. Acad. Sci. 118, e2026099118 (2021).

59. F. Masina, R. Pezzetta, S. Lago, D. Mantini, C. Scarpazza, G. Arcara, Disconnection from prediction: A systematic review on the role of right temporoparietal junction in aberrant predictive processing. Neurosci. Biobehav. Rev. 138, 104713 (2022).

60. A. Ali, N. Ahmad, E. de Groot, M. A. J. van Gerven, T. C. Kietzmann, Predictive coding is a consequence of energy efficiency in recurrent neural networks (2021), p. 2021.02.16.430904,, doi:10.1101/2021.02.16.430904.

61. A. HajiHosseini, A. Rodríguez-Fornells, J. Marco-Pallarés, The role of beta-gamma oscillations in unexpected rewards processing. NeuroImage. 60, 1678–1685 (2012).

62. S. Lloyd-Fox, A. Blasi, G. Pasco, T. Gliga, E. J. H. Jones, D. G. M. Murphy, C. E. Elwell, T. Charman, M. H. Johnson, Cortical responses before 6 months of life associate with later autism. Eur. J. Neurosci. 47, 736–749 (2018).

63. M. F. Siddiqui, C. Elwell, M. H. Johnson, Mitochondrial Dysfunction in Autism Spectrum Disorders. Autism-Open Access. 6, 1000190 (2016).

64. G. Bale, S. Mitra, I. de Roever, M. Sokolska, D. Price, A. Bainbridge, R. Gunny, C. Uria-Avellanal, G. S. Kendall, J. Meek, N. J. Robertson, I. Tachtsidis, Oxygen dependency of mitochondrial metabolism indicates outcome of newborn brain injury. J. Cereb. Blood Flow Metab. (2018), p. 0271678X1877792.

65. A. Vezyroglou, P. Hebden, I. De Roever, R. Thornton, S. Mitra, A. Worley, M. Alves, E. Dean, J. H. Cross, I. Tachtsidis, Broadband-NIRS System Identifies Epileptic Focus in a Child with Focal Cortical Dysplasia&mdash;A Case Study. Metabolites. 12 (2022),, doi:10.3390/metabo12030260.

66. M. Jeong, M. Tashiro, L. N. Singh, K. Yamaguchi, E. Horikawa, M. Miyake, S. Watanuki, R. Iwata, H. Fukuda, Y. Takahashi, M. Itoh, Functional brain mapping of actual car-driving using [18F]FDG-PET. Ann. Nucl. Med. 20, 623–628 (2006).

67. I. Lundgaard, B. Li, L. Xie, H. Kang, S. Sanggaard, J. D. R. Haswell, W. Sun, S. Goldman, S. Blekot, M. Nielsen, T. Takano, R. Deane, M. Nedergaard, Direct neuronal glucose uptake heralds activity-dependent increases in cerebral metabolism. Nat. Commun. 6, 6807 (2015).

68. A. B. Rocher, F. Chapon, X. Blaizot, J.-C. Baron, C. Chavoix, Resting-state brain glucose utilization as measured by PET is directly related to regional synaptophysin levels: a study in baboons. NeuroImage. 20, 1894–1898 (2003).

69. S. N. Vaishnavi, A. G. Vlassenko, M. M. Rundle, A. Z. Snyder, M. A. Mintun, M. E. Raichle, Regional aerobic glycolysis in the human brain. Proc. Natl. Acad. Sci. U. S. A. (2010), doi:10.1073/pnas.1010459107.

70. S. Lloyd-Fox, A. Blasi, A. Volein, S. Lloyd-Fox, A. Blasi, A. Volein, N. Everdell, C. E. Elwell, M. H. Johnson, Social perception in infancy: a near infrared spectroscopy study. Child Dev. (2009), doi:10.1111/j.1467-8624.2009.01312.x.

71. S. Lloyd-Fox, M. Papademetriou, M. K. Darboe, N. L. Everdell, R. Wegmuller, A. M. Prentice, S. E. Moore, C. E. Elwell, Functional near infrared spectroscopy (fNIRS) to assess cognitive function in infants in rural Africa. Sci. Rep. 4, 4740 (2014).

72. P. Phan, D. Highton, J. Lai, M. Smith, C. Elwell, I. Tachtsidis, Multi-channel multi-distance broadband near-infrared spectroscopy system to measure the spatial response of cellular oxygen metabolism and tissue oxygenation. Biomed. Opt. EXPRESS. 7 (2016), doi:10.1364/BOE.7.004424.

73. B. Molavi, G. A. Dumont, Wavelet-based motion artifact removal for functional near-infrared spectroscopy. Physiol Meas Physiol Meas. 33, 259–270 (2012).

74. A. Duncan, J. Meek, M. Clemence, C. Elwell, L. Tyszczuk, M. Cope, D. Delpy, Optical pathlength measurements on adult head, calf and forearm and the head of the newborn infant using phase resolved optical spectroscopy. Phys. Med. Biol. 40, 295 (1995).

75. J. M. Kilner, J. Mattout, R. Henson, K. J. Friston, Hemodynamic correlates of EEG: A heuristic. NeuroImage (2005), doi:10.1016/j.neuroimage.2005.06.008.

76. M. J. Rosa, J. Kilner, F. Blankenburg, O. Josephs, W. Penny, Estimating the transfer function from neuronal activity to BOLD using simultaneous EEG-fMRI. NeuroImage (2010), doi:10.1016/j.neuroimage.2009.09.011.

77. Y. Benjamini, Y. Hochberg, Controlling the false discovery rate: a practical and powerful approach to multiple testing. J. R. Stat. Soc. B. 57, 289–300 (1995).

78. K. J. Friston, A. P. Holmes, K. J. Worsley, J.-P. Poline, C. D. Frith, R. S. J. Frackowiak, Statistical parametric maps in functional imaging: A general linear approach. Hum. Brain Mapp. 2, 189–210 (1994).

79. M. L. Schroeter, M. M. Bücheler, K. Müller, K. Uludağ, H. Obrig, G. Lohmann, M. Tittgemeyer, A. Villringer, D. Y. von Cramon, Towards a standard analysis for functional near-infrared imaging. NeuroImage. 21, 283–290 (2004).

80. S. Shimada, K. Hiraki, Infant’s brain responses to live and televised action. NeuroImage. 32, 930–939 (2006).

81. Y. Minagawa-Kawai, H. van der Lely, F. Ramus, Y. Sato, R. Mazuka, E. Dupoux, Optical Brain Imaging Reveals General Auditory and Language-Specific Processing in Early Infant Development. Cereb. Cortex. 21, 254–261 (2011).

82. D. Arifler, T. Zhu, S. Madaan, I. Tachtsidis, Optimal wavelength combinations for near-infrared spectroscopic monitoring of changes in brain tissue hemoglobin and cytochrome c oxidase concentrations. Biomed. Opt. Express (2015), doi:10.1364/boe.6.000933.

83. F. Shi, P. T. Yap, G. Wu, H. Jia, J. H. Gilmore, W. Lin, D. Shen, Infant brain atlases from neonates to 1- and 2-year-olds. PLoS ONE (2011), doi:10.1371/journal.pone.0018746.

84. M. Jenkinson, M. Pechaud, S. Smith, “BET2: MR-based estimation of brain, skull and scalp surfaces” in Eleventh Annual Meeting of the Organization for Human Brain Mapping (2005).

85. S. Brigadoi, P. Phan, D. Highton, S. Powell, R. J. Cooper, J. Hebden, M. Smith, I. Tachtsidis, C. E. Elwell, A. P. Gibson, Image reconstruction of oxidized cerebral cytochrome C oxidase changes from broadband near-infrared spectroscopy data. Neurophotonics (2017), doi:10.1117/1.NPh.4.2.021105.

86. A. Corlu, R. Choe, T. Durduran, K. Lee, M. Schweiger, S. R. Arridge, E. M. C. Hillman, A. G. Yodh, Diffuse optical tomography with spectral constraints and wavelength optimization. Appl. Opt. (2005), doi:10.1364/AO.44.002082.

87. M. Schweiger, S. Arridge, The Toast++ software suite for forward and inverse modeling in optical tomography. J. Biomed. Opt. (2014), doi:10.1117/1.jbo.19.4.040801.

88. F. Bevilacqua, D. Piguet, P. Marquet, J. D. Gross, B. J. Tromberg, C. Depeursinge, In vivo local determination of tissue optical properties: applications to human brain. Appl. Opt. (1999), doi:10.1364/ao.38.004939.

89. G. Strangman, J. P. Culver, J. H. Thompson, D. A. Boas, A quantitative comparison of simultaneous BOLD fMRI and NIRS recordings during functional brain activation. NeuroImage. 17, 719–731 (2002).

90. A. Custo, W. M. Wells, A. H. Barnett, E. M. C. Hillman, D. A. Boas, Effective scattering coefficient of the cerebral spinal fluid in adult head models for diffuse optical imaging. Appl. Opt. (2006), doi:10.1364/AO.45.004747.

91. J. Zhao, H. S. Ding, X. L. Hou, C. Le Zhou, B. Chance, In vivo determination of the optical properties of infant brain using frequency-domain near-infrared spectroscopy. J. Biomed. Opt. (2005), doi:10.1117/1.1891345.

92. M. A. Franceschini, S. Thaker, G. Themelis, K. K. Krishnamoorthy, H. Bortfeld, S. G. Diamond, D. A. Boas, K. Arvin, P. E. Grant, Assessment of Infant Brain Development With Frequency-Domain Near-Infrared Spectroscopy. Pediatr. Res. (2007), doi:10.1203/pdr.0b013e318045be99.

