## Supplementary information for "Mapping human social brain specialisation beyond the neuron using multimodal imaging in human infants"

**Supplementary Material**

### 1 Social and non-social

#### 1.1 EEG results

Statistical tests were performed to compare the social and non-social conditions to the baseline and to compare the social condition to the non-social condition. The results from these are shown in Tables 1, 2 and 3.

| **Social versus baseline** | | | |
| --- | --- | --- | --- |
| Theta | Alpha | Beta | Gamma |
| C3 (p-value = 0.0416, t-value = 2.229)  P10 (p-value = 0.0133, t-value = 2.805)  PO8 (p-value = 0.0437, t-value = 2.2021) | C4 (p-value = 0.0313, t-value = 2.438) | P3 (p-value = 0.0172, t-value = 2.679)  C4 (p-value = 0.0211, t-value = 2.651)  F4 (p-value = 0.0455, t-value = 2.428)  FC6 (p-value = 0.0113, t-value = 3.036) | P7 (p-value = 0.0232, t-value = 2.527)  **Cz (p-value = 0.0133, t-value = 2.807)***  FC6 (p-value = 0.0494, t-value = 2.208)  C2 (p-value = 0.0494, t-value = 2.1513)  TP7 (p-value = 0.0283, t-value = 2.428)  FC5 (p-value = 0.0471, t-value = 2.212) |

Table 1: Statistical tests performed on average RMS power in the stimulus period versus average RMS power in the baseline period for the social condition. The p-values and t-values are shown in brackets and bold values indicated with an asterisk indicate the channels significant after FDR correction.

| **Non-social versus baseline** | | | |
| --- | --- | --- | --- |
| Theta | Alpha | Beta | Gamma |
| C3 (p-value = 0.0431, t-value = -2.210)  FC6 (p-value = 0.0437, t-value = -2.307) | FC6 (p-value = 0.0127, t-value = -3.031) | P7 (p-value = 0.0410, t-value = -2.212)  P3 (p-value = 0.037, t-value = -2.263)  O1 (p-value = 0.048, t-value = -2.131)  C3 (p-value = 0.0274, t-value = -2.443)  PO7 (p-value = 0.0287, t-value = -2.390)  CP5 (p-value = 0.039, t-value = -2.236)  P9 (p-value = 0.0237, t-value = -2.50) | Pz (p-value = 0.0216, t-value = -2.529)  O1 (p-value = 0.0176, t-value = -2.630) |

Table 2: Statistical tests performed on average RMS power in the stimulus period versus average RMS power in the baseline period for the non-social condition. The p-values and t-values are shown in brackets and bold values indicated with an asterisk indicate the channels significant after FDR correction.

| **Social versus Non-social** | | | |
| --- | --- | --- | --- |
| Theta | Alpha | Beta | Gamma |
| **C3 (p-value = 0.0041, t-value = 3.108)***  FC6 (p-value = 0.0232, t-value = 2.448) | FC6 (p-value = 0.0371, t-value = 2.225) | **P3 (p-value = 0.011, t-value = 2.70)***  C3 (p-value = 0.032, t-value = 2.246) | **Pz (p-value = 0.002, t-value = 3.375)***  P3 (p-value = 0.047, t-value = 2.0654)  F8 (p-value = 0.034, t-value = 2.275)  **C4 (p-value = 0.0096, t-value = 2.824)***  FC6 (p-value = 0.045, t-value = 2.130) |

Table 3: Statistical tests performed on average RMS power in the stimulus period for the social condition versus the non-social condition. The p-values and t-values are shown in brackets and bold values indicated with an asterisk indicate the channels significant after FDR correction.

Figure 1 shows the scalp topographies of the RMS power in each 1 s segment of the block consisting of 1 s of pre-stimulus onset and 6 s post-stimulus onset (for visualisation purposes) for each of the frequency bands for the social condition, the non-social condition and the social minus the non-social condition.

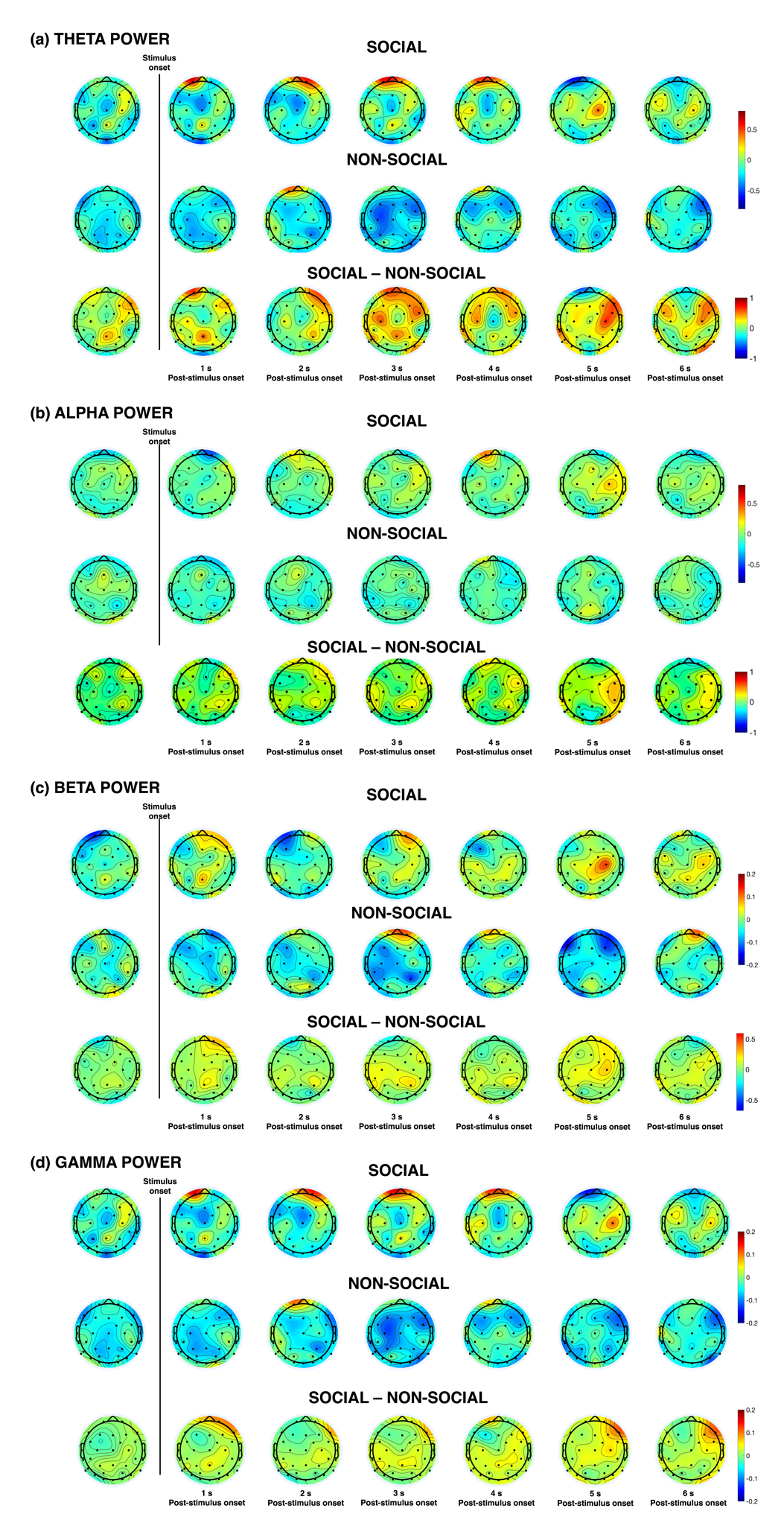

Figure 1: Scalp topographies of the RMS power block for (a) theta, (b) alpha, (c) beta, and (d) gamma frequency bands for the social condition, the non-social condition and social minus non-social condition.

#### 1.2 Combined bNIRS-EEG results

The β-values that were estimated for each bNIRS and EEG channel combination from the GLM analysis were averaged within hemispheres. For the social condition, the highest average β-value of within the right hemisphere was observed for the gamma band for HbO_2_ (0.195) and HHb (0.062) and for the beta band for oxCCO (0.027).

Tables 4 and 5 show the significant bNIRS and EEG connections. The p-values are shown and t-values are shown in brackets.

| **Frequency band** | **EEG channel** | **bNIRS** | | | | | |
| --- | --- | --- | --- | --- | --- | --- | --- |
|  |  | **HbO2** | | **HHb** | | **oxCCO** | |
|  |  | Channel 12 | Channel 14 | Channel 12 | Channel 14 | Channel 12 | Channel 14 |
| **Theta band** | O2 | - | - | 0.013  (2.812) | - | - | - |
|  | P10 | - | - | 0.016  (2.723) | - | - | - |
| **Alpha band** | - | - | - | - | - | - | - |
|  | - | - | - | - | - | - | - |
| **Beta band** | Pz | - | 0.039 (2.295) | 0.048  (-2.180) | - | - | 0.032  (2.40) |
|  | TP8 | 0.003  (-3.624) | - | - | - | - | - |
|  | T8 | 0.006  (-3.334) | - | - | - | - | - |
| **Gamma band** | Pz | - | 0.0043  (2.529) | - | - | - | - |
|  | PO4 | - | 0.027  (2.501) | - | - | - | 0.025  (2.529) |

Table 4: Statistical values of connections between bNIRS and EEG channels for HbO_2_, HHb and oxCCO for the social condition, for all frequency bands**.** p-values are shown, and t-values are shown in brackets.

| **Frequency band** | **EEG channel** | **bNIRS** | | |
| --- | --- | --- | --- | --- |
|  |  | **HbO2** | **HHb** | **oxCCO** |
|  |  | Channel  14 | Channel  14 | Channel  14 |
| **Theta band** | P4 | - | - | 0.008  (3.051) |
|  | P10 | - | - | 0.006  (3.244) |
|  | PO8 | - | - | 0.01  (2.999) |
|  | O2 | - | - | 0.006  (3.219) |
| **Alpha band** | P10 | - | - | 0.002  (3.843) |
| **Beta band** | CP6 | 0.0114  (-2.859) | - | - |
|  | C2 | - | 0.005  (-3.280) | 0.002  (3.827) |
|  | PO8 | - | 0.006  (-3.248) | 0.000018  (6.344) |
|  | O2 | - | 0.023  (-2.534) | 0.003  (3.515) |
| **Gamma band** | P4 | - | - | 0.009  (2.96) |
|  | Cz | - | 0.0146  (-2.738) | - |
|  | C2 | - | 0.004  (-3.325) |  |
|  | Oz | - | 0.016  (-2.769) | - |

Table 5: Statistical values of connections between bNIRS and EEG channels for HbO_2_, HHb and oxCCO for the non-social condition, for all frequency bands**.** p-values are shown, and t-values are shown in brackets.
